# Fruit flies subjected to TBI exhibit genotype-dependent changes in seizure incidence and duration

**DOI:** 10.1101/2021.10.26.465914

**Authors:** Tori R Krcmarik, Ashley M Willes, A Yang, Sydney L Weber, Douglas J Brusich

## Abstract

Traumatic brain injury (TBI) is common and often debilitating. One complication following TBI is post-traumatic seizures (PTS). However, there is a poor understanding of PTS, in part, because it is challenging to model. We sought to develop a fly model of PTS. We used a high-impact trauma (HIT) device to inflict TBI and followed this with behavioral, bang-testing methods to assess seizure behavior. Our results showed PTS incidence was sensitive to genetic background. We also found seizure duration was most pronounced at 30 minutes after injury. Our findings support the efficacy of our fly model for coarse screening of seizure behavior. We expect this model will be useful in first-pass screens intended to identify modifiers of seizure risk following TBI.

## Introduction

Traumatic brain injury (TBI) is a common occurrence, which accounts for millions of cases each year globally (Majdan et al. 2016; CDC: Centers for Disease Control 2016). TBI can give rise to multiple complications, which may include ataxia and seizures (Mysiw et al. 1990; Agrawal et al. 2006). Seizure conditions following TBI are estimated to account for 5% of all epilepsies (Agrawal et al. 2006). Seizures which occur within the first week of injury are termed post-traumatic seizures (PTS), and most early seizures occur within the first 24 hours following injury (Jennett 1973; Agrawal et al. 2006; Verellen and Cavazos 2010). The overall rate of early PTS correlates with the severity of the primary injury (Annegers et al. 1998; Verellen and Cavazos 2010; Najafi et al. 2015). Early seizures are also predictive for late and recurrent seizures, termed post-traumatic epilepsy (PTE) (Jennett 1973; Annegers et al. 1998; Englander et al. 2003; Verellen and Cavazos 2010). However, the underlying genetic predisposition and molecular factors that link TBI to either PTS or PTE are poorly understood.

Fruit flies (*Drosophila melanogaster*) hold promise as a useful model for studying PTS and PTE. Fly seizure behavior can be evaluated at multiple levels, including behaviorally, genetically, and electrophysiologically (Lee and Wu 2002; Song and Tanouye 2008; Iyengar and Wu 2014; Stone et al. 2014). More recently, flies have become established as a TBI model and primary injuries are replicable across varying levels of injury severity (Katzenberger et al. 2013; Rebeccah J. Katzenberger et al. 2015; Rebeccah J Katzenberger et al. 2015; Putnam et al. 2019; Willes et al. 2021). Moreover, the genetic and molecular tool kits available to the fly model make flies a well-suited choice for downstream analysis of the basis of PTS and PTE.

In this experiment, we developed and evaluated a simple fly model of PTS. We hypothesized flies would exhibit increased seizure activity following TBI dependent upon time post-injury and injury condition. To test this hypothesis, we used flies of two commonly used genetic backgrounds, *w^1118^* and *y^1^w^1^*. Flies were injured using three separate levels of TBI severity and evaluated for seizure behavior at three time points post-injury. We found *y^1^w^1^* flies experienced greater seizure incidence at both 30 minutes and 2 hours post-injury for each of the three injury conditions relative to uninjured controls. Moreover, after injury by each of the three conditions used, *y^1^w^1^* flies exhibited longer seizure durations at 0.5h post-injury than for events evoked in uninjured flies. By contrast, *w^1118^* flies showed no statistical change in seizure incidence nor duration for any condition relative to uninjured controls. Our results demonstrate utility for this fly model of PTS, such as in a preliminary screen for genetic modifiers of PTS incidence. Our data also provides a starting-point for additional validation and detailed seizure characterization by electrophysiological techniques.

## Materials and Methods

### Fly husbandry and TBI methodology

The Bloomington Drosophila Stock Center (Bloomington, Indiana, USA) was the source for both the *w^1118^* (BL 5905) and *y^1^w^1^* (BL 1495) fly stocks. A 25°C humidified incubator set on a 12H:12H light:dark cycle was used for housing flies. Flies were kept on a glucose-cornmeal-yeast media (Putnam et al. 2019), and were collected for TBI experiments using light CO_2_ anesthesia by 5 days after eclosion (dae).

Flies were hand-transferred from media-containing vials to empty test vials in preparation for TBI. TBI was administered to the test vials at various injury severities using the HIT device, as done previously (Putnam et al. 2019). Injury conditions included in this study were 70° x 3HITs, 80° x 2HITs, and 90° x 1HIT. Uninjured controls were handled identically, but were not given TBI.

### Bang-testing methodology

Injured and uninjured flies were hand-transferred back to media-containing vials post-injury and allowed to recover in the incubator. Flies were then assessed for seizure behavior at 0.5hr, 2hr, or 24hr post-injury using bang-testing methods. Separate cohorts of flies were used for each time-point. To prepare flies for bang-testing, they were hand-transferred to an empty vial, anesthetized on ice for 5 minutes, and transferred to new empty vials at a density of approximately 5-6 flies per vial. Flies were given 10 minutes in vials at room temperature for recovery, and then vortexed for 10 seconds at the maximum setting to evoke seizure activity (Vortex Genie 2). Behavior was recorded using a webcam (Logitech c270) until all flies showed a full recovery, or up to a maximum of 60 seconds after termination of vortex treatment.

### Video analysis

Seizure videos were replayed and scored for seizure behavior (VLC Mediaplayer, version 3.0.6). Flies that experienced convulsive behavior, which included shaking, spinning, and wing-buzzing were noted as exhibiting seizure behavior. Start and end times were determined for each fly’s first seizure. Flies that continued to seize up to the 60 second mark were capped at 60 seconds duration. The seizure duration was calculated by subtracting the seizure start-time from the seizure end-time. Only event times for the first seizure event for each fly were used in the analysis.

### Statistics

Comparisons of categorical count data (seizure:no seizure) between injury conditions were conducted using a 2 x 2 Fisher’s Exact Test at α = 0.01. Statistical testing of seizure durations between uninjured *w^1118^* and *y^1^w^1^* flies was done via a Mann Whitney U-test at α = 0.05. Statistical testing of seizure durations within genotypes was done via Kruskal Wallis testing with multiple comparisons of mean ranks and Dunn’s correction at α = 0.05. All statistical testing and visualization of data was done using GraphPad Prism 7.05 (GraphPad Software, San Diego, California USA).

## Results

### Seizure incidence analysis

Cohorts of flies were injured by one of three separate injury conditions, 70° x 3HITs, 80° x 2HITs, or 90° x 1HIT. These conditions were previously shown to result in comparable levels of mortality at 24hr post-injury (Putnam et al. 2019). However, the conditions vary by primary injury severity, with 90° causing the most severe injuries and 70° the least severe injuries.

Flies were assessed for seizures post-injury at either 0.5hr, 2hr, or 24hr by bang-testing, which entailed 10 seconds of vortex treatment followed by observation of fly behavior. We noted that many flies showed brief locomotive changes immediately after vortex treatment, including in the uninjured condition (22/92 uninjured *w^1118^* flies = 23.9%; 40/146 uninjured *y^1^w^1^* flies = 27.4%). However, not all of these changes showed evidence of sustained loss of coordination or robust seizure activity. In order to exclude ordinary, non-seizure events, we elected to only include events which lasted for greater than 1.0s in our analysis of seizure behaviors, which is consistent with the multiple seconds-long seizure events characterized previously (Lee and Wu 2002).

The number of flies that exhibited seizure behavior vs. those that did not were compared across genotypes and injury conditions. We found no differences in seizure incidence between uninjured genotypes (Fig. 1A). For comparisons across TBI conditions, we found *w^1118^* flies were generally resistant to changes in seizure incidence compared to their uninjured controls (Fig. 1B). None of the injury conditions in *w^1118^* flies exhibited changes in seizure incidence vs. uninjured controls (Fig. 1B). We did, however, note an approximately 62% reduction in seizure incidence in the 80° x 2HITs condition from 0.5hr to 2hr post-injury (Fig. 1B: 21.5% seizures at 80° x 2HITs and 0.5hr (45/209) vs. 8.2% seizures at 2hr post-injury (8/97)). By contrast, injury resulted in increased seizure incidence in *y^1^w^1^* flies compared to uninjured controls across all conditions except 24hr post-injury time-points for 80° x 2HITs and 90° x 1HIT (Fig. 1C). Statistically significant increases in seizure incidence in *y^1^w^1^* flies ranged from 2.2- to 3.2-fold (80° x 2HITs at 2hr vs uninjured controls and 90° x 1HIT at 0.5hr vs uninjured controls respectively)

**Figure 1.**
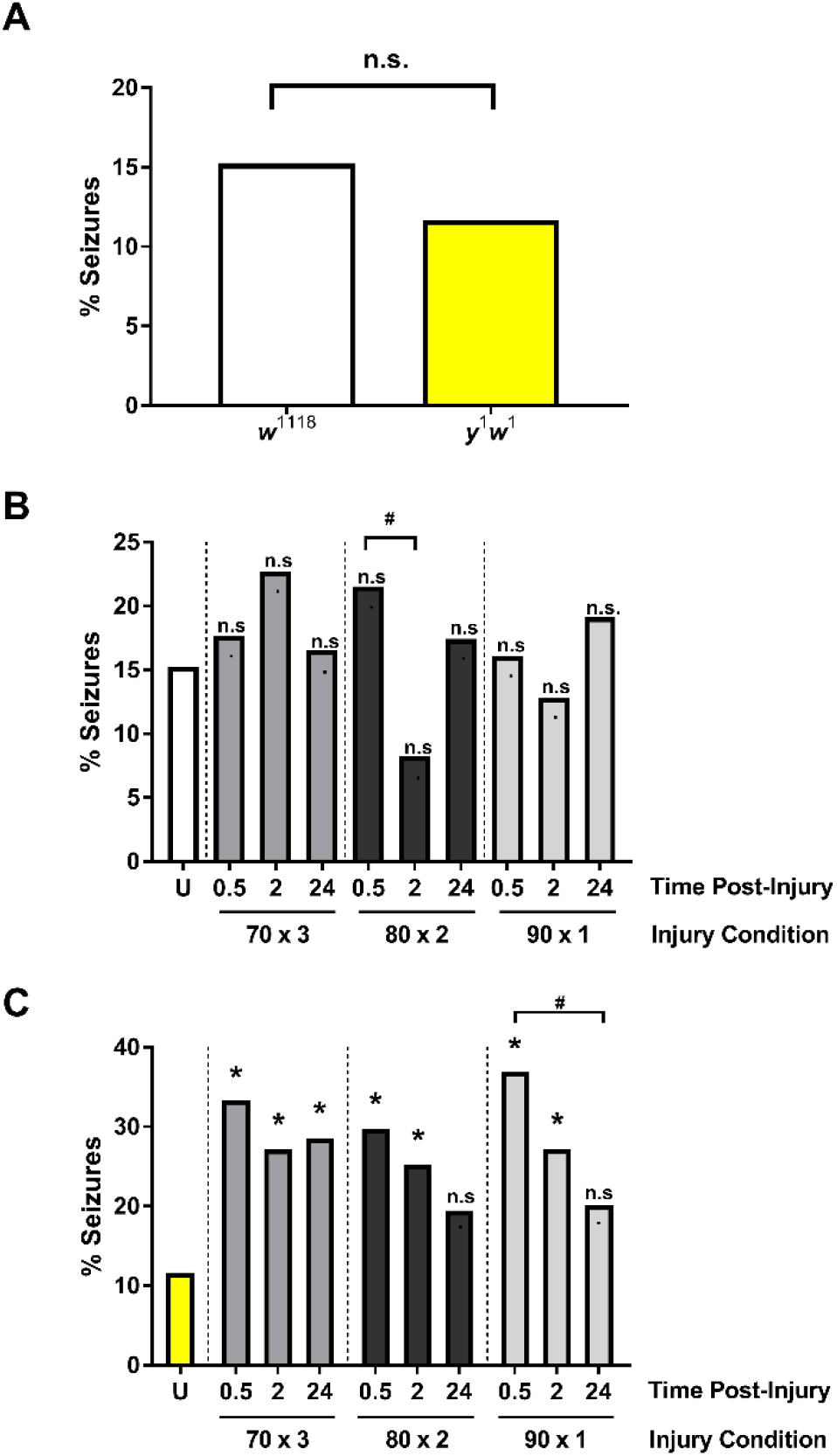
*y^1^w^1^* flies, but not *w^1118^* flies, exhibit increased seizure incidence post-injury. (A) Seizure incidence in uninjured *w^1118^* and *y^1^w^1^* flies was not different. Seizure incidence was determined across three injury conditions at three time-points each in both (B) *w^1118^* and (C) *y^1^w^1^* flies. Only *y^1^w^1^* flies exhibited increased seizure incidence post-injury compared to uninjured controls (U). n.s.= p > 0.05 between indicated conditions or relative to uninjured controls (U). * = p < 0.01 for indicated condition vs. respective uninjured controls (U). # = p < 0.01 between indicated conditions. All statistical testing via Fisher’s Exact Test. n ≥ 78 flies for each condition for *w^1118^*. n ≥ 45 flies for each condition for *y^1^w^1^*.

### Seizure duration analysis

In addition to seizure incidence, we measured seizure durations from the start of locomotor dysfunction to its resolution. We did not see a difference in seizure durations between uninjured genotypes (Fig. 2A). Further, there were no changes in seizure duration for *w^1118^* flies across injury conditions compared to uninjured controls (Fig. 2B). However, we did note that injured *w^1118^* flies trended towards longer seizure durations at 0.5hr post-injury. Within *y^1^w^1^* flies we found a robust increase in seizure duration for all three injury conditions when assessed at 0.5hr post-injury (Fig. 2C). At this 0.5hr time-point the median seizure duration was increased from 2.8-fold (90° x 1HIT) to 5.9-fold (70° x 3HITs). The *y^1^w^1^* flies also showed longer seizure times within the 70° x 3HITs injury condition at 0.5hr post-injury compared to each of the 2hr and 24hr post-injury times (Fig. 2C).

**Figure 2.**
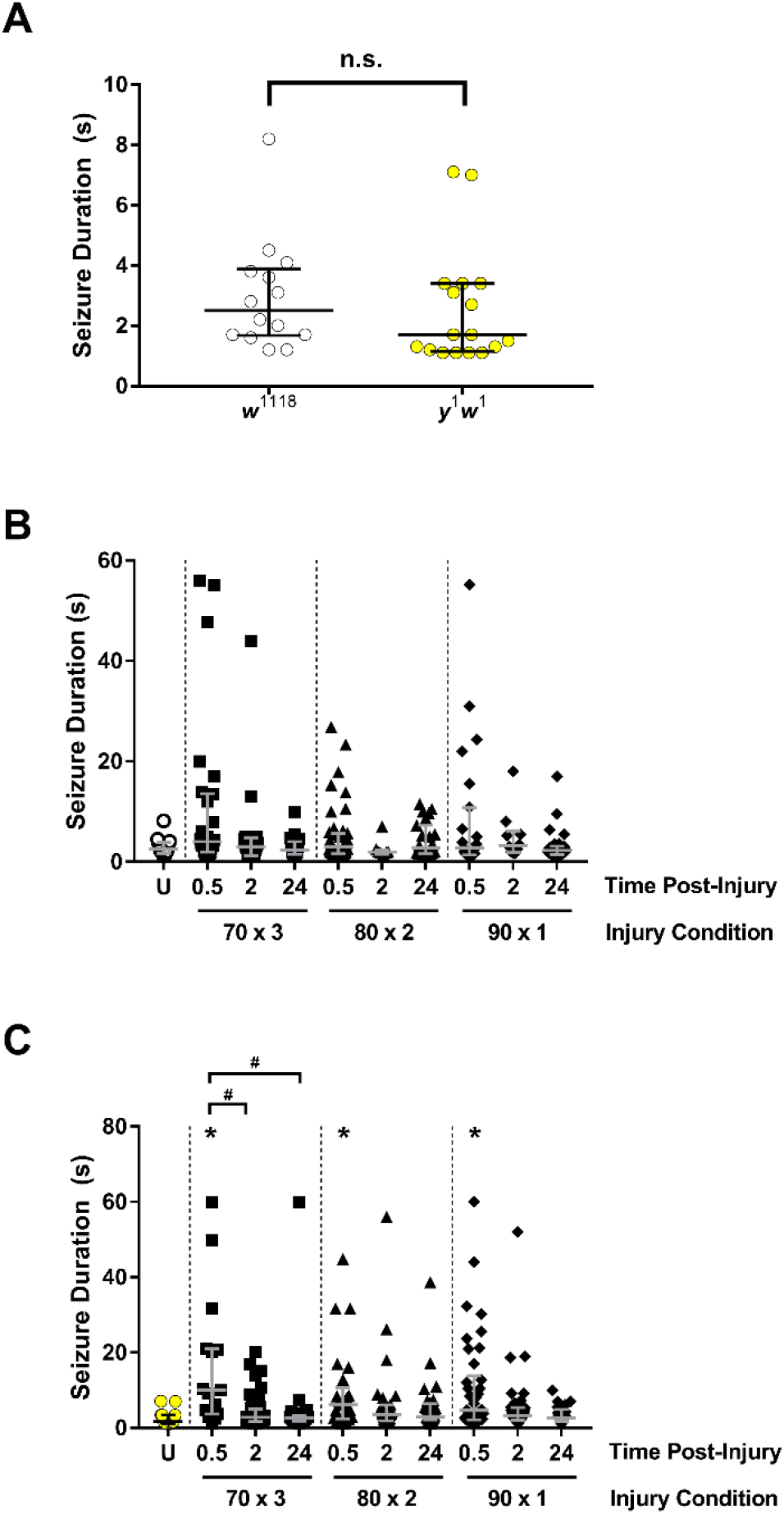
*y^1^w^1^* flies show longer seizures at 0.5hr relative to controls, while *w^1118^* flies trend towards the same pattern. (A) Seizure durations were no different in uninjured controls. n.s.= p > 0.05 by Mann-Whitney U-test. n ≥ 14 seizure events per condition. Seizure durations across injury conditions were compared for (B) *w^1118^* flies and (C) *y^1^w^1^* flies. * = p < 0.05 vs. uninjured control and # = p < 0.05 between indicated conditions via Kruskal Wallis with Dunn’s Correction. n≥ 8 seizure events per condition for *w^1118^*. n ≥15 seizure events for *y^1^w^1^*. Each symbol represents the first seizure event from a single fly. Error bars indicate the median and IQR.

## Discussion

Our study revealed two main findings. First, seizure incidence was dependent upon genotype, with *y^1^w^1^* flies, but not *w^1118^* flies, exhibiting increased seizure incidence post-injury (Fig. 1). Second, we found longer seizure durations at 0.5hr post-injury in *y^1^w^1^* flies compared to controls, and a trend towards this same relationship in *w^1118^* flies (Fig. 2).

The genetic bases for PTS or PTE prevalence are poorly understood and only a few genetic risk factors have been identified. One gene associated with PTS is *GAD1*, which encodes an enzyme necessary for synthesis for the inhibitory neurotransmitter gamma-Aminobutyric acid (GABA) (Darrah et al. 2013). This and other evidence implicate inhibitory-excitatory balance as important in PTS, but validation of target factors involved in this process and a model amenable to testing such factors remain problematic (Guerriero et al. 2015). The PTS model described in this study and the evidence that seizure incidence is sensitive to genetic background will be useful in addressing this gap.

In humans, most seizures following TBI occur within the first 24hr (Jennett 1973; Agrawal et al. 2006). In this study, we found that the 0.5hr post-injury time point was best for detecting seizure activity, and seizures at this time point tended to be longer and more severe (Fig. 2). In humans, early seizures within the first 24hr are more common in children than adults, and children are more likely to suffer longer lasting status epilepticus seizures (Jennett 1973). We used young flies (≤ 5 dae) in our experiment. It is an open question whether or not older flies would show differences in seizure activity compared to age-matched controls when injured.

Overall, our model holds promise for further study of PTS. The flexibility of the fly system and genetic tools allow for systematic genetic screening and validation of genetic factors that predispose animals to PTS. Further, this system allows testing of multiple time points post-injury, multiple injury conditions, and varied ages which allow for a more comprehensive model. One drawback of our design is the lack of objective measure of seizure threshold such as by electroconvulsive seizure (ECS) analysis (Lee and Wu 2002). However, our approach offers a simple first pass analysis of PTS activity that can be confirmed and further characterized with more in-depth approaches such as ECS.

## Author Contributions

<u>Tori R Krcmarik</u>: Formal Analysis, Writing – Original Draft, Writing – Review and Editing, Visualization

<u>Ashley M Willes</u>: Investigation, Writing – Review and Editing

<u>A Yang</u>: Investigation, Writing – Review and Editing

<u>Sydney L Weber</u>: Investigation, Writing – Review and Editing

<u>Douglas J Brusich</u>: Formal Analysis, Investigation, Writing – Review and Editing, Conceptualization, Methodology, Project administration, Resources, Supervision, Visualization

